# MRI Guided Fresh Tissue Procurement in Radical Prostatectomy Specimens: An Evolutionary Paradigm

**DOI:** 10.1101/2022.05.09.491189

**Authors:** Raiza Z Freidlin, Jonathan Krynitsky, Cody Bastian, Connor Schultz, Julian Custer, Sarah Young, Sherif Mehralivand, Maria J Merino, Vladimir A Valera, Peter Pinto, Baris Turkbey, Choyke L Peter, Thomas J Pohida, Marcial A Garmendia

## Abstract

**Purpose:** The aim of this work is to assess the feasibility of novel targeted methods for fresh tissue procurement that facilitate accurate identification of lesion location and tissue sample extraction in an excised prostate gland, while minimizing the tissue deformation or damage.

**Methods and Results:** Proposed fresh tissue procurement methods are based on utilizing a custom 3D printed patient-specific prostate mold. In the semi-freehand method, the access cut is guided by a set of markers on the bottom of the mold that define the boundary of a lesion in two orthogonal directions. The procurement site is identified by palpation. In the guided cut with-locator card method the procurement sites are identified by a cutout in the locator cards inserted into the access cuts. The two approaches were utilized to procure >50 samples. In the biopsy needle method guiding channels were designed into the prostate mold to procure fresh tissue. This method was used 4 times. Finally, a mixed-reality biopsy needle method was developed for phantom testing.

**Conclusions:** Preliminary results of fresh tissue procurement utilizing novel methods showed improve precision of obtaining cancerous fresh tissue for molecular and genomic cancer research compared to current methods which rely on surgical estimation of lesion location.

## 1. Introduction

Prostate cancer is the most common non-cutaneous cancer type, and one of the leading causes of cancer death in men living in the United States [1]. Comprehensive research of prostate cancer at the molecular level provides the basis for improved diagnosis, prognosis, and treatment planning and often require procurement of fresh tissue from recently excised specimens. Typically, metabolomic profiling, gene expression, and protein pattern analyses are performed to assess molecular level changes associated with cancer [2,3]. Therefore, there is a continuous need for more effective fresh tissue procurement methods of prostate cancers. Currently, the surgeon estimates the location of the cancer within the prostate and excises a section of tissue from the specimen. Unfortunately, the actual cancer is often missed with this approach. A more accurate approach would be welcomed provided it did not interfere with subsequent histopathological clinical evaluations.

Radical prostatectomy specimens are the ideal source of fresh tissue for genetic and metabolic studies. However, targeted fresh tissue procurement from cancer foci within the resected prostate glands has its own challenges. In particular, after prostatectomy, the *in vivo* orientation and shape of the gland is largely lost as the surrounding supporting tissues are no longer present and this makes identification of precise lesion location more difficult.

In order to address this problem, several different techniques have been developed. Wheeler et al. harvested 5mm long cylindrical samples of fresh tissue from a single access cut through the mid portion of the prostate by performing approximately 10 random punch biopsies [4]. The non-targeted nature of this approach leads to a large number of unproductive samples. Walton et al. utilized a custom designed needle to perform at least 8 biopsies in a standard sextant scheme with an additional two lateral biopsies within the specimen [5]. However, as in the previous method, this non-targeted approach also has a low yield of relevant samples. A number of procurement techniques involve slicing the entire unfixed prostate gland and locating cancerous tissue prior to procurement [6,7]. This requires a skilled pathologist as the soft consistency of the fresh prostate makes it hard to cut accurately and is subject to inter-operator variability. Moreover, this procurement damages the entire specimen making clinical assessment more difficult. Another set of existing techniques are based on sectioning of the gland and blind harvesting of the tissue [8,9,10], which have the disadvantages associated with non-targeted sampling.

In this study, we assess the feasibility of novel targeted methods for fresh tissue procurement that facilitate accurate lesion localization and tissue sample extraction in an excised gland. Given prior MRI-based knowledge of the lesion location, these techniques utilize either a single partial incision or needle biopsies, both of which are conducted with the gland secured and oriented by a patient-specific prostate mold [11]. Such evolutionary approaches aim to minimize the tissue deformation, misalignment, or damage caused by slicing of the entire gland that may hinder further histopathology clinical reads or imaging validation studies.

## 2. Methods and Results

A critical component of the prostate fresh tissue procurement process is designing a multi-parametric MRI-based (mpMRI) patient-specific mold [11], which is based on a computer-rendered 3D model of the prostate gland derived from T2 weighted MRI. The computer-rendered prostate 3D model is generated from the boundaries of the prostate gland and the cancer foci are outlined by a trained radiologist on three orthogonal planes of the mpMRI. The patient-specific mold establishes spatial correlation between the excised prostate gland and mpMRI imaging by preserving the *in-vivo* orientation and shape of the organ, thus enabling accurate identification of cancer lesion location(s) within the excised gland. The mold assembly consists of two parts, designated the “top” and “bottom”. The patient specific mold has been used for correlations studies of imaging and histopathology in our center since 2008 and over 1000 molds have been made [11]. In addition, since January 2017 we have utilized the mold for fresh tissue procurement. The evolution of our approach to procurement has undergone three different generations representing continuous improvement in the accuracy and quality of the procured tissue.

### Semi-freehand cut method

The semi-freehand method was the first iteration of our customized mold assisted procurement technique. This method utilizes the boundary points of a lesion along two orthogonal directions. An experienced radiologist outlines the edges of a prostate lesion generating a 3D lesion model. The markers are defined based on the location of the 3D lesion model embedded within the prostate 3D model and are annotated via two sets of opposing markers on either the top or bottom of the mold (Figure 1). It is important to note that there are no indicators to define the depth of the access cut. The surgeon performs a freehand cut of the specimen based on the markers with the cut depth at the discretion of the surgeon (Figure 2b). Tissue procurement is assisted by palpation of the cancer by an experienced surgeon (Figure 2d).

**Figure 1.**
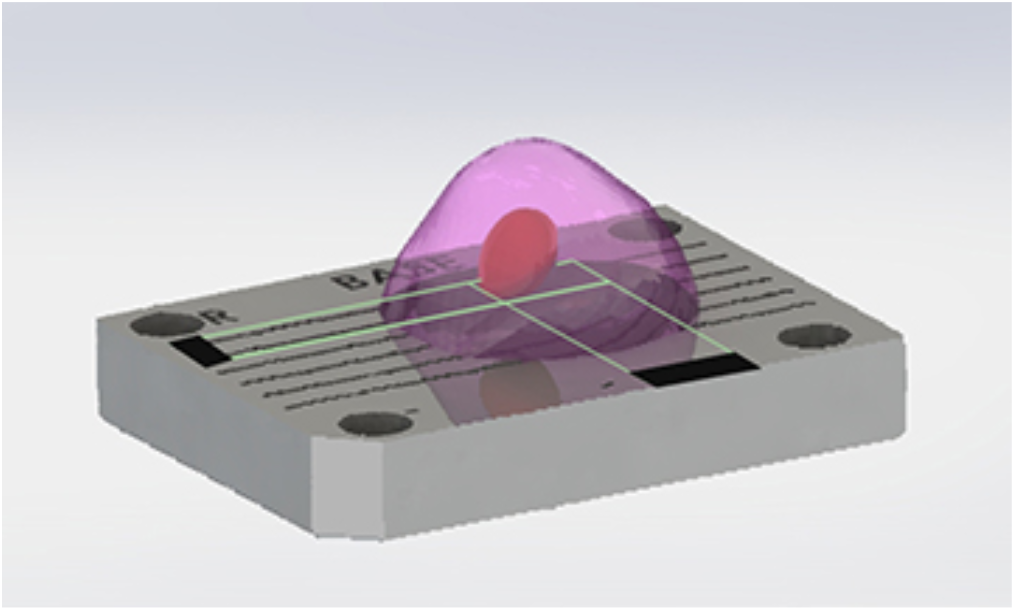
Schematic drawing of a mold bottom with outlined boundaries of a lesion and corresponding notched markers for rough placement of access cut.

**Figure 2.**
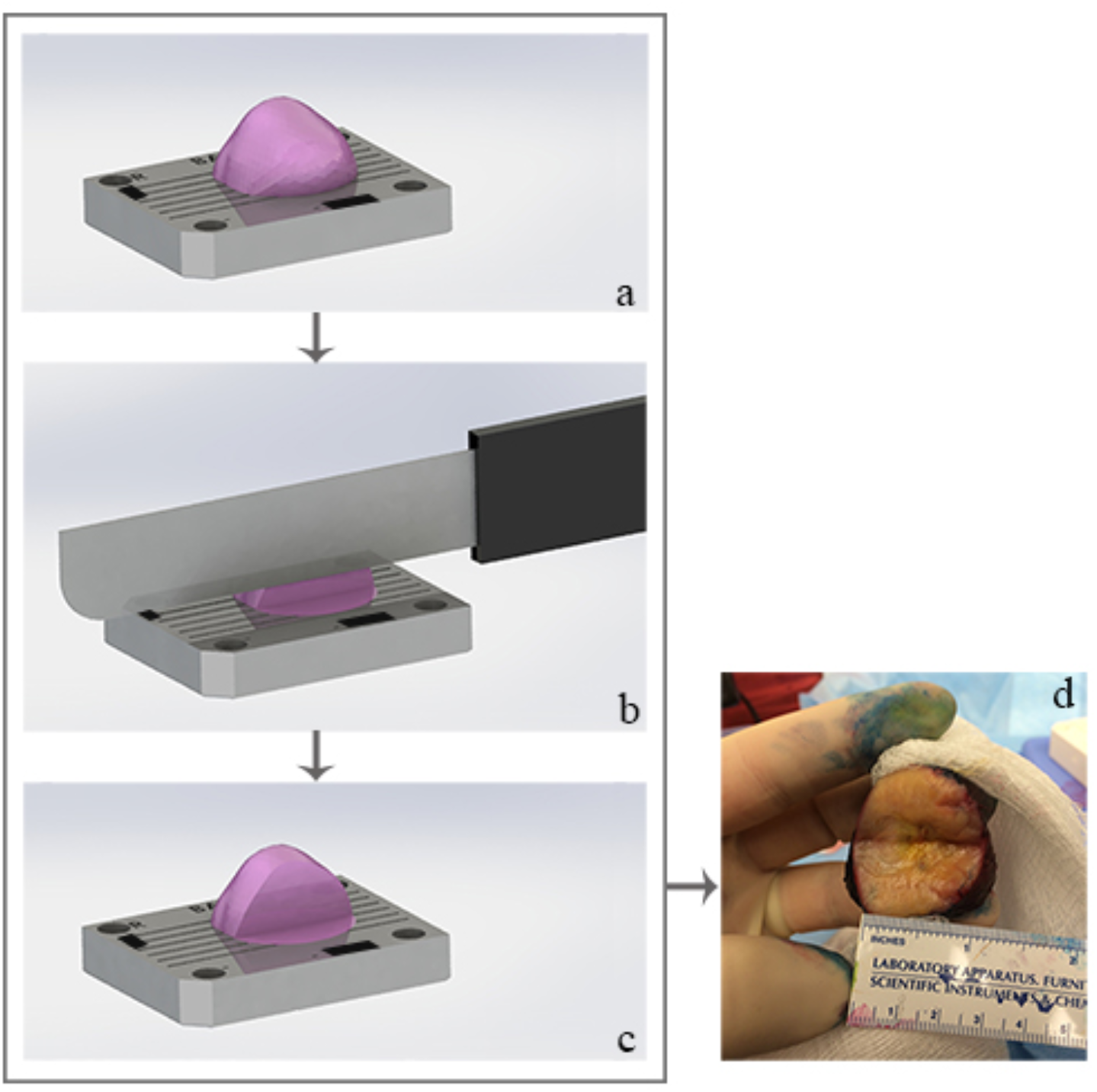
Process of semi-freehand prostate cancer tissue procurement consists of placing (a) excised prostate into the bottom part of a patient-specific sectioning-mold. (b) The freehand cut is coincident with the markers, with the cut depth at the discretion of the surgeon. (c) Access cut. (d) Excised prostate after the procurement access cut.

Semi-freehand cut method was utilized in 6 prostatectomies for procurement over 2 months.

### Guided cut with locator card method

Although the semi-freehand cut technique improves tissue procurement localization, the method is still subject to errors due to potential subjectivity of the cut depth and relies on palpation to determine the depth of the cut which can be misleading. To address the depth-of-cut uncertainty, we developed an improved method using both parts of the split prostate mold using an integrated procurement slot to guide a knife through the prostate lesion, stopping the cut at a predetermined depth (Figure 3a and 3b). Once the cut is made a lesion “locator card” further defines the lesion location (Figure 3c). The shape of the locator card is determined by the profile of the segmented prostate gland in the plane of the access cut. The cutout within the card corresponds to the manual and planimetric tumor segmentation performed by an experienced radiologist. The locator card is inserted into the access cut and a tumor tissue sample is obtained through the card cutout as shown in Figure 3d.

**Figure 3.**
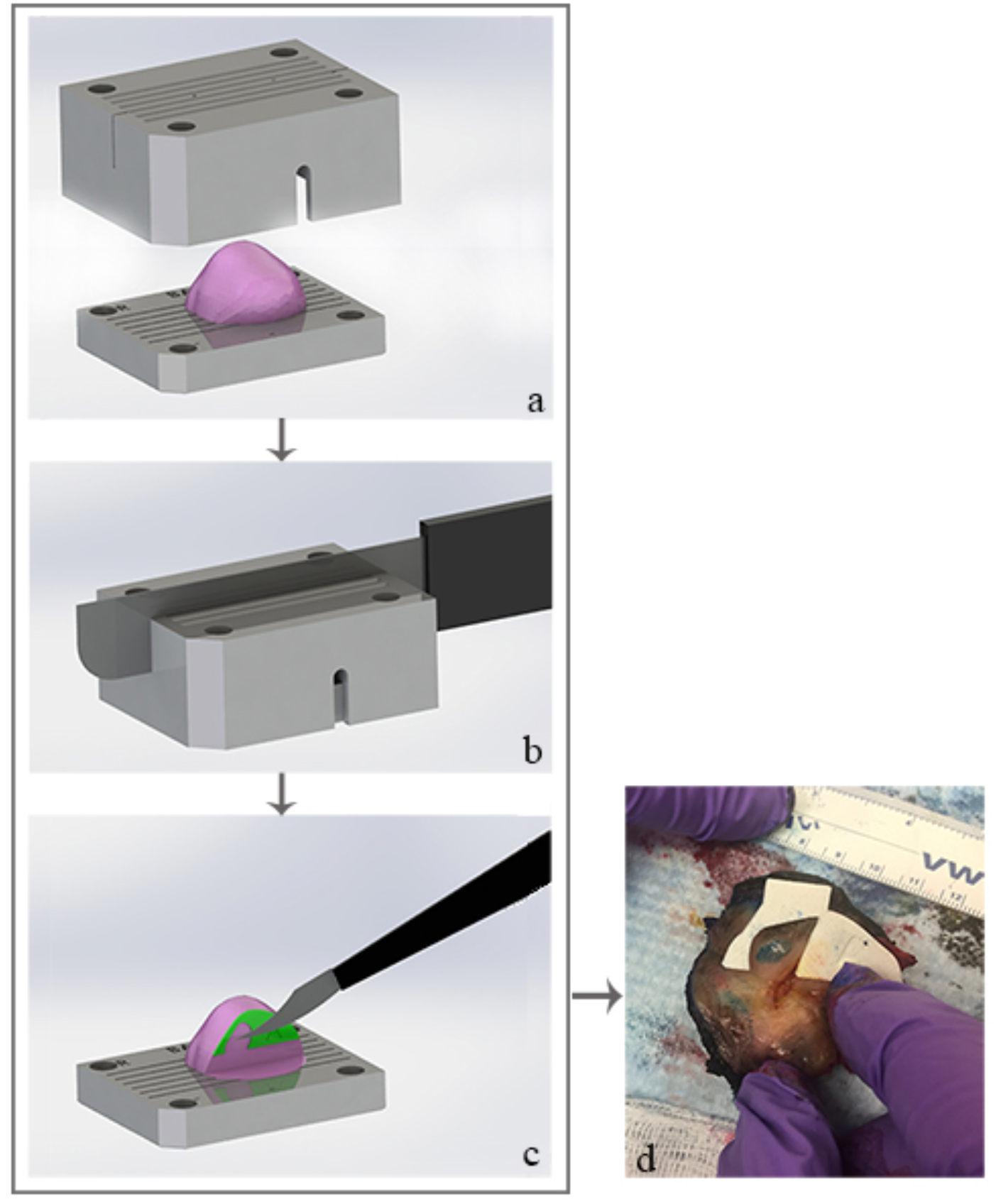
Process of precision prostate cancer tissue procurement consists of placing (a) excised prostate into the patient-specific sectioning-mold. (b) The lesion access cut knife slot is incorporated into the mold and the cut depth is limited to the lesion depth. (c) A locator card with an outline of the lesion (green) is placed in the procurement cut for fresh tissue procurement. (d) Excised prostate with a locator card placed in the procurement cut.

Figure 4a shows histology slide with procurement area within the left anterior mid based and a corresponding axial T2-weighted MR image with lesion location identified by an experienced radiologist (Figure 4b).

**Figure 4.**
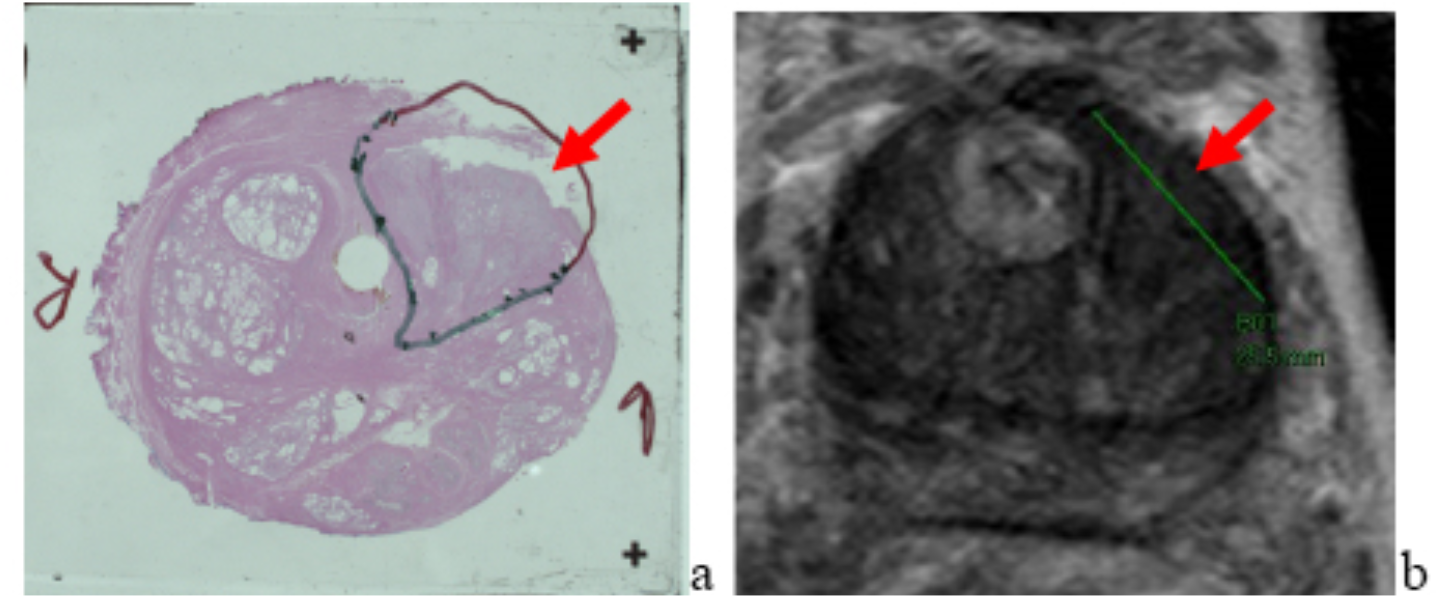
(a) Histology slide with a location of tissue procured from the left midbase anterior transition zone prostate cancer lesion. (b) Corresponding axial T2-weighted MR image with lesion location.

### Guided biopsy needle procurement method

One of the limitations of cutting the prostate gland with a knife is potential deformation of the organ. In particular, the previously described access-cut methods can potentially hinder subsequent clinical interpretation of the specimen and further studies that require correlation between histopathology slides and corresponding *in-vivo* mpMRI slices. To reduce the tissue disturbance, we implemented a less intrusive approach utilizing biopsy needles. For this purpose, instead of guiding knife cuts, we integrated needle guiding channels into the patient-specific prostate mold. To ensure a sufficient volume of tissue collection, multiple (e.g., 3) needle guiding channels are designed into the mold for each lesion. The depth of needle insertion is controlled by the length of a custom sleeve placed over the needle. The sleeve is specific to each channel. Figure 5 schematically shows the procurement procedure performed with a biopsy needle and guide channels within in the customized mold.

**Figure 5.**
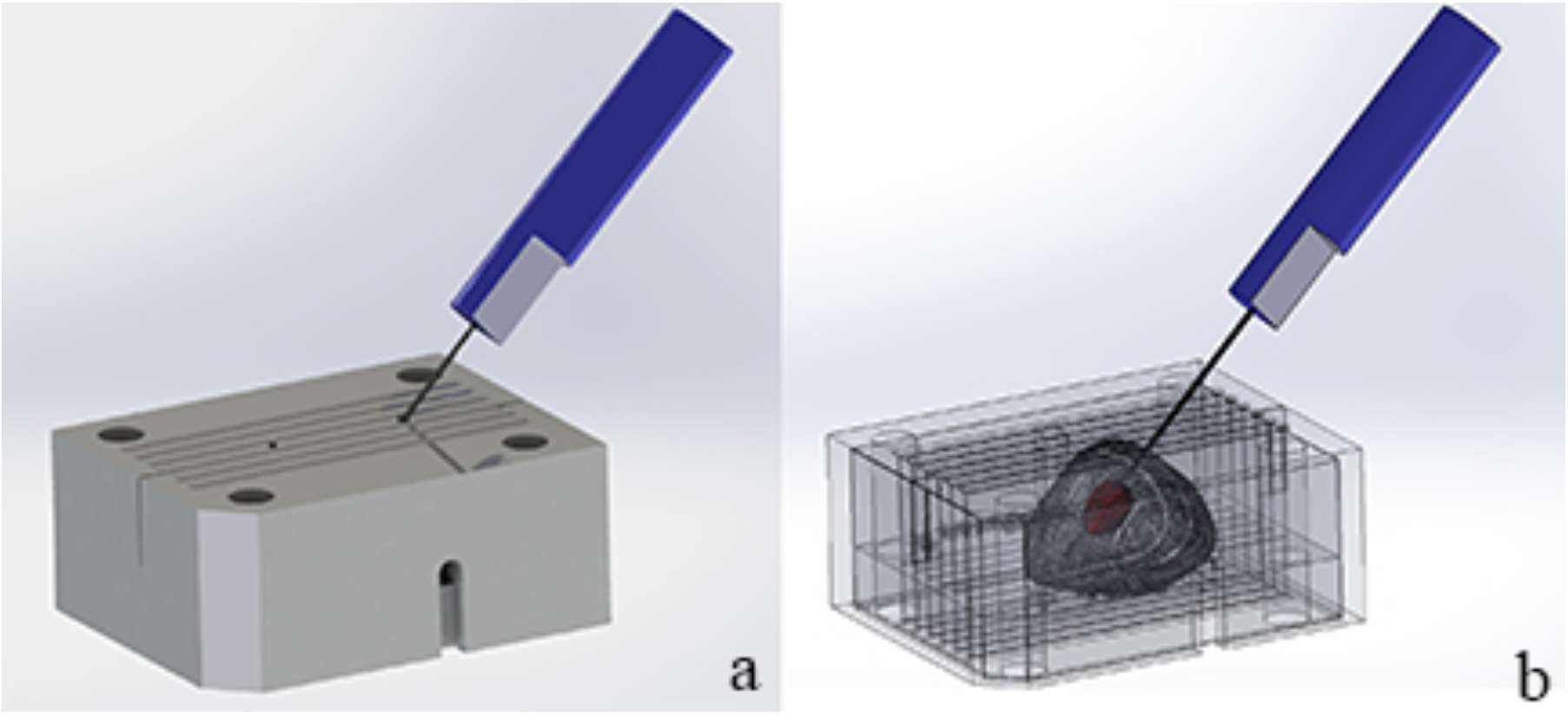
(a) Prostate mold with the guiding channels. (b) Wireframe rendering of the procedure with biopsy needle penetrating into an intraprostatic lesion.

### Mixed reality guided biopsy needle procurement method

In the previously described fresh tissue procurement methods, the prostate gland is enclosed within the customized mold, thus procurement is performed without visual verification of where biopsy needle enters the gland. Additionally, the number and path of mold-guided biopsies are limited due to mechanical challenges of incorporating additional guide channels. To address these challenges, we have recently proposed a next generation method which is still under development (Figure 6). This novel method aims to guide a biopsy needle into prostate cancer lesion for tissue procurement with assistance of Mixed Reality (MR). An ensemble of computer-generated 3D models of a prostate gland and embedded lesion is overlaid on the actual gland, placed in a mold bottom part, and displayed in MR goggles. The real-world biopsy needle is tracked and represented as a hologram in MR goggles, thus allowing flexibility selecting the insertion path by visualizing the location of the needle within the gland. Furthermore, the lack of in-mold constraints resolves limitation on number of biopsies that can be performed.

**Figure 6.**
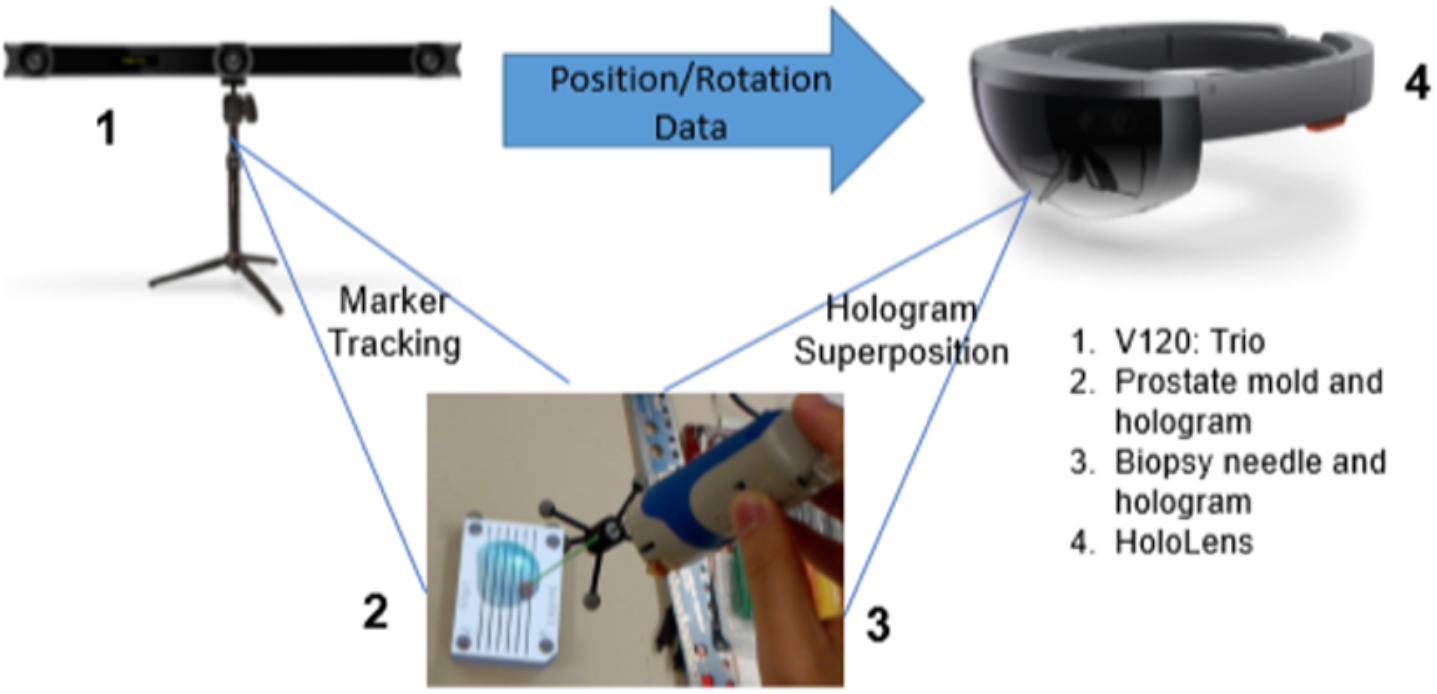
Schematic overview of the mixed reality guided procurement system setup.

## 3. Discussion

Fresh tissue procurement is an essential aspect of basic and translational oncology research. Fresh tissue is needed for many advanced proteomic and genetic analyses as formalin fixed tissue often degrades the desired signal. However, there are a number of challenges associated with obtaining abundant fresh samples, such as accurate localization within the tumor while not disturbing the specimen in a way that makes routine clinical interpretation difficult. To meet these challenges, we propose four novel fresh tissue procurement methods which evolved as our experience grew. The first two methods require cuts within the prostate gland in order to reach the tumor tissue, whereas the latter two methods enable users to obtain tissue via biopsy needles without specimen incision. The first two approaches were utilized in >50 samples, in which fresh tumor tissue was harvested successfully, whereas the channel guided biopsy needle procurement method enabled us to obtain fresh tumor tissue in 4 prostatectomy samples. An advantage of these techniques, compared to the existing non-targeted methods, is utilization of the mold to restore the *in-vivo* orientation and shape of the organ, which, along with lesion segmentation, allows for intuitive localization of a lesion for targeted procurement. In addition, all four methods minimize organ destruction during tissue procurement. Due to the semi-automated mold design process and reduced printing time (only mold top or bottom is required), this method is practical to implement and offers significant localization information to improve procurement.

Although the semi-freehand cut is the simplest to implement, the final location for the procurement is identified through palpation, which is highly subjective. The second approach, guided cut with locator card, allows precise extraction of the cancerous sample. However, these first two methods require incision into the prostate to extract samples. This causes partial organ destruction, thus hindering consecutive histopathology studies and clinical evaluation. The guided biopsy needle procurement techniques, channel guided and utilizing mixed reality, significantly reduce damage and deformation of the gland. The channel guided method presents its own challenges, such as the number and angle of the channels (i.e., samples) due to mold manufacturing process and lack of visual monitoring of potential needle bending. Mixed reality biopsy needle guiding improves flexibility of sample extraction and provides real-time visual feedback. However, it still requires rigorous testing with real prostatectomy samples.

## 4. Conclusions

The fresh tissue procurement procedures, using novel approaches with utilization of the patientspecific prostate mold, improve precision of tissue acquisition for molecular and genomic cancer research. Future work is required to further determine the actual impact of these novel procurement methods on performance of genomic and proteomic research efforts.

## Supporting information

Prostate Mold Automated Generation Process

## Notes

### Competing Interest Statement

The authors have declared no competing interest.

