## Supplementary material for "MRI Guided Fresh Tissue Procurement in Radical Prostatectomy Specimens: An Evolutionary Paradigm": Prostate Mold Automated Generation Process: Prostate mold automation process.pdf

**Process flow diagram:**

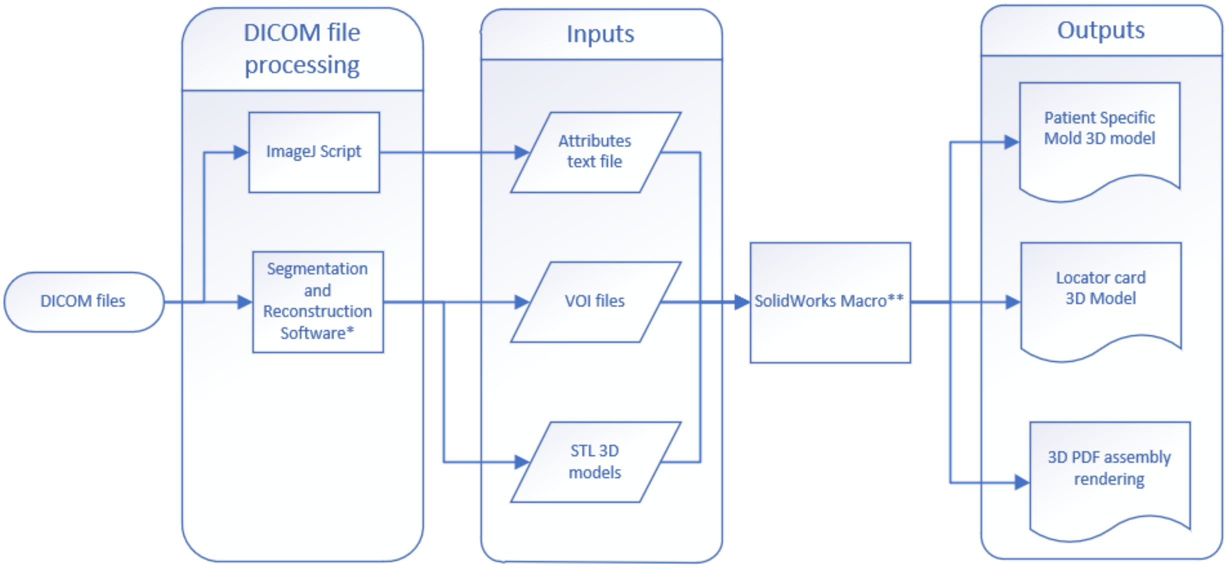


*Provided by Radiologist(s). Segmentation and reconstruction can be separate software.
**Step-by-step instructions found below

The SolidWorks Application Programming Interface (API) tool is used to create a custom program (i.e., macro) to assist in automating the design of the patient-specific prostate mold used for fresh tissue procurement and gland blocking to 6mm slices prior to fixation. The input data for the macro is extracted from the DICOM images obtained from MRI or CT scans. List of files:

1. Inputs
2. SolidWorks macro for mold generation
3. Outputs

### 1. Inputs

1.1. Attributes text file (DICOM metadata)
DICOM metadata contains a series of attributes that identifies image parameters such as dimensions, pixel resolution, and slice orientation in the world coordinate system. An ImageJ script extracts these attributes from the DICOM file and generates a text file (DICOM.txt) used by the SolidWorks macro to establish the world coordinate system. Step-by-step instructions can be found in “**Mold generation –1. Inputs**: 1.1 DICOM input” section.

1.2. Contouring information in image space (SLICER VOI file format)
Trained radiologist(s) segment/outline regions of interest (ROIs), such as organs/lesions, from stacks of two-dimensional (2D) MRI images, to generate a vector file (*.VOI) that contains contouring information in 2D image space. The SolidWorks macro parses through the axial

*.VOI file to extract data values such as the total number of slices and the vertices of the ROIs in the corresponding slices. The extrema coordinates along the X and Y directions and the location of the first and last slices that contain ROIs along the Z direction are used to calculate the outline of the mold (Figure 1) by translating 2D coordinates from the image space to the world coordinate system. The location of the sectioning slots is determined by the first slice and spacing (i.e., 6mm) between slices. Step-by-step instructions can be found in "**Mold generation – 1. Inputs:** 1.2 VOI input" section.

1.3. Computer-rendered 3D models (STL file)
A prostate 3D model is reconstructed from the ROIs segmented in three orthographic views and saved as *.STL files. The SolidWorks macro imports the STL 3D models and orients them relative to the world coordinate system. The macro uses the 3D model of the prostate to generate the mold cavity, while the 3D model of the lesion(s) is used to generate the procurement features in the mold and locator card. The use of a urethra 3D model to improve prostate orientation and stability within the mold is optional. Step-by-step instructions are in "**Mold generation – 1. Inputs:** 1.3 STL input" section.

### 2. SolidWorks macro for mold generation summary (step-by-step instructions start on page 10)

The SolidWorks macro automates the mold generation process using a Graphical User Interface (GUI). Main functions:

2.1 Calculation of the optimal parting line (Figure 2) resulting in the 2-piece mold.
2.2 Reconciliation of the different medical imaging coordinate systems into the world coordinate system.
2.3 Creation of 3D printable files.
2.4 Generation of assembly files (Figure 3) and an interactive 3D PDF of the entire assembly for tissue procurement and gland blocking procedure planning.

### 3. Outputs

3.1. Patient-specific mold 3D model (STL format)
Top and bottom parts of the 2-piece mold are separated along the coronal plane that contains the largest prostate profile. A "lesion-access cut" is incorporated into the mold based on the lesion location.

3.2. Locator card 3D model (STL format)
The perimeter shape of the locator card is determined by the profile of the segmented prostate gland in the plane of the lesion-access cut. The outline (i.e., void) of the lesion within the card is determined by the profile of the segmented lesion in the lesion-access cut, which allows for a targeted tissue procurement.

3.3. 3D PDF assembly rendering (PDF format)

The interactive (e.g., zoom, pan, rotate) assembly rendering includes the patient-specific mold, procurement locator card, prostate, lesions, and urethra (optional) 3D models.

Figures:

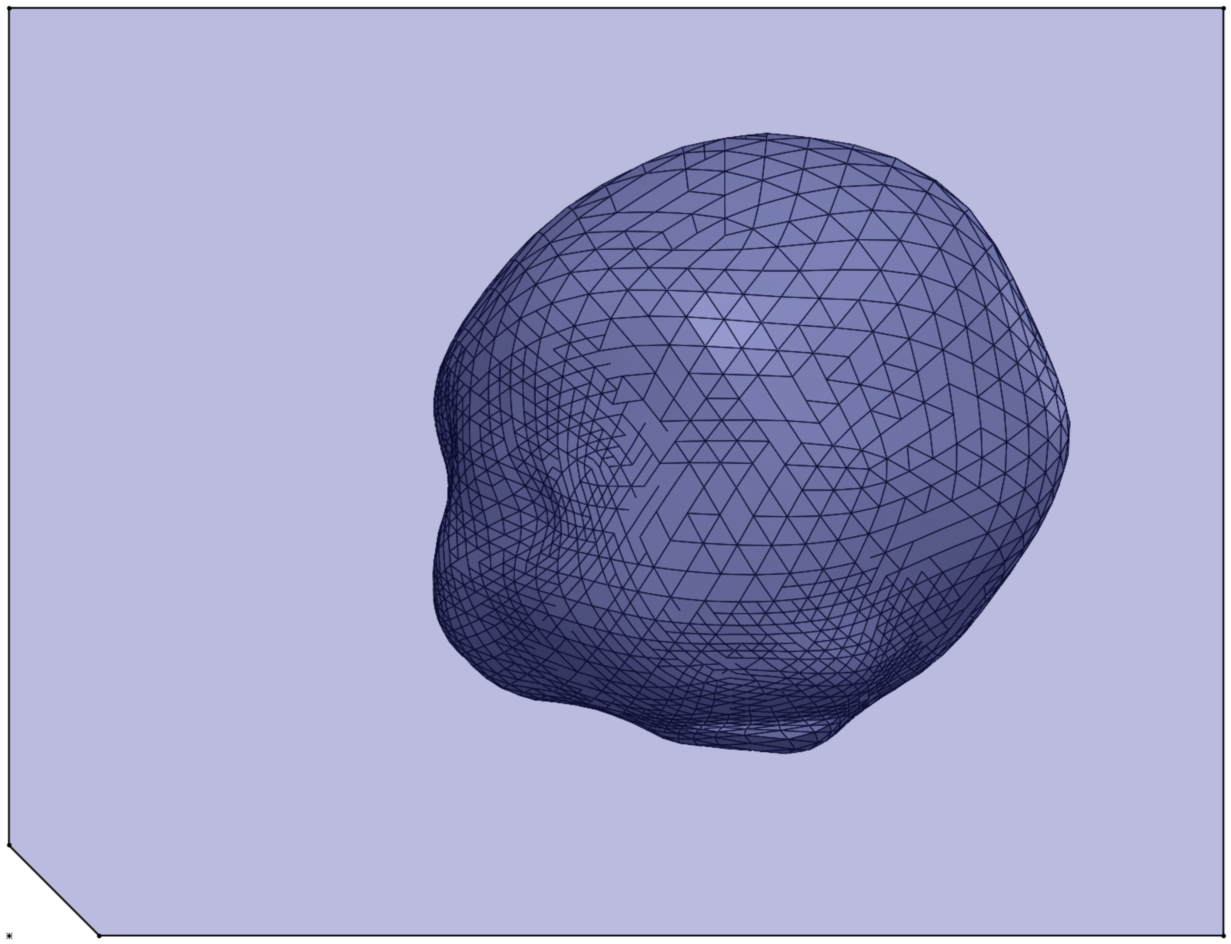

*Figure 1- Outline of the mold generated around the prostate gland.*

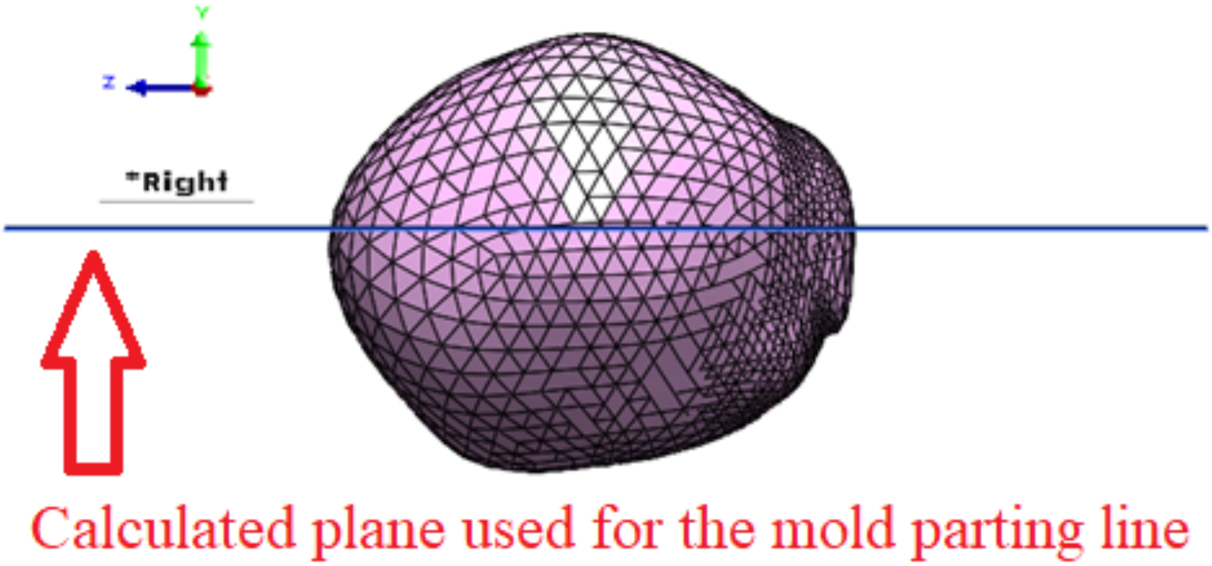


*Figure 2 – The mold parting plane (the X-Z plane) is shown in blue*

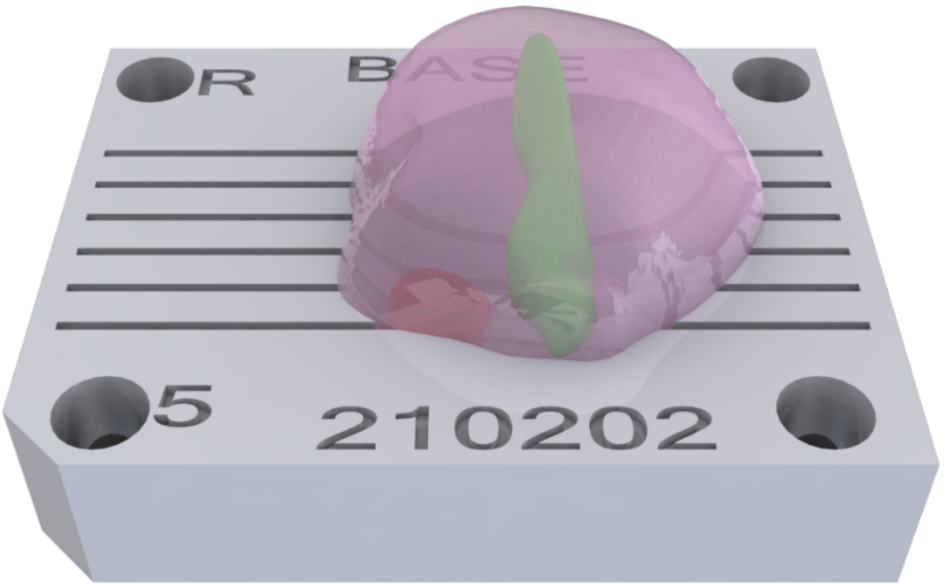


*Figure 3 - SolidWorks mold assembly: bottom part of the mold (gray), prostate (purple), tumor (red), and urethra (green)*

### Mold Generation

1. **Inputs:**

1.1. <u>DICOM Input</u>
Software required: ImageJ
Required script: ImageJ script: "MB_GetDICOM4MOLDSW.txt"

**Step 1: Create the "patientmolds" folder on your C: drive, and add "axial" and "mold" sub-folders.**

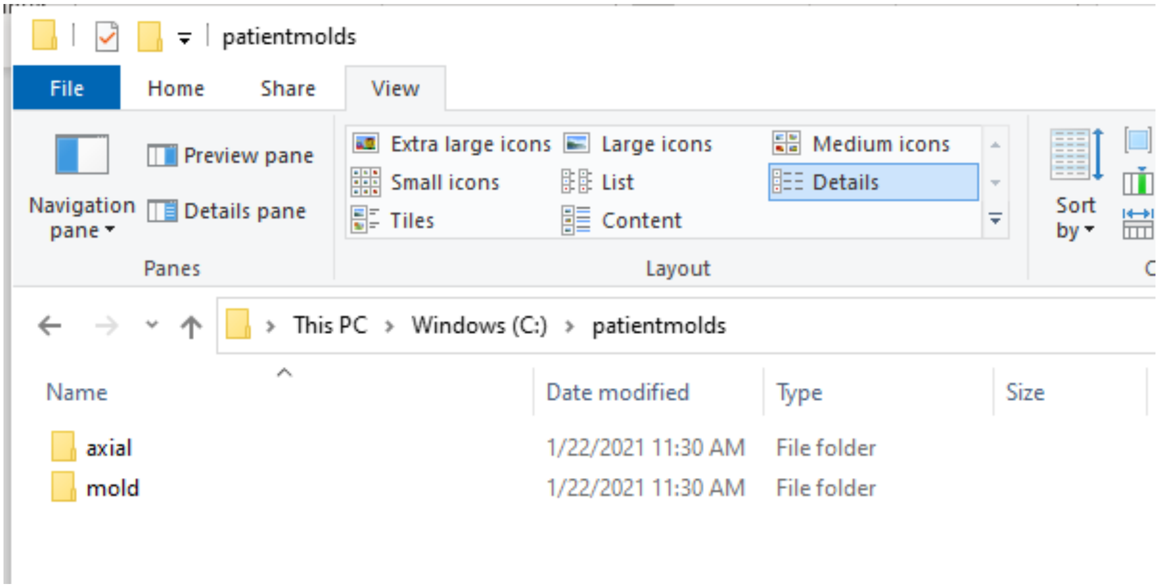


**Step 2: Copy the ImageJ script "MB_GetDICOM4MOLDSW.txt" to the "patientsmold/axial" folder.**

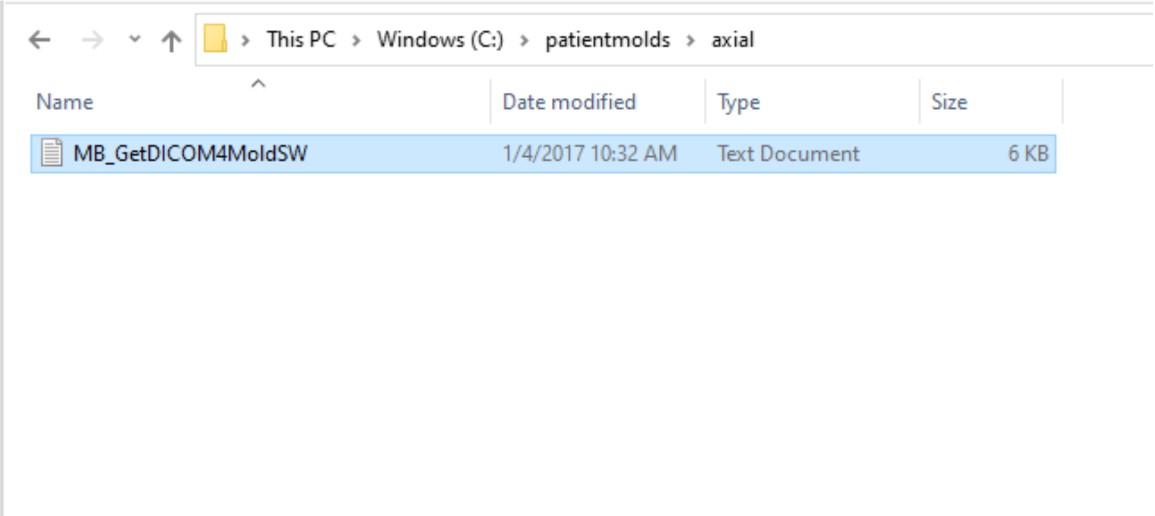


**Step 3: Copy the entire stack of axial DICOM images to the "C:\patientsmold\axial" folder.**

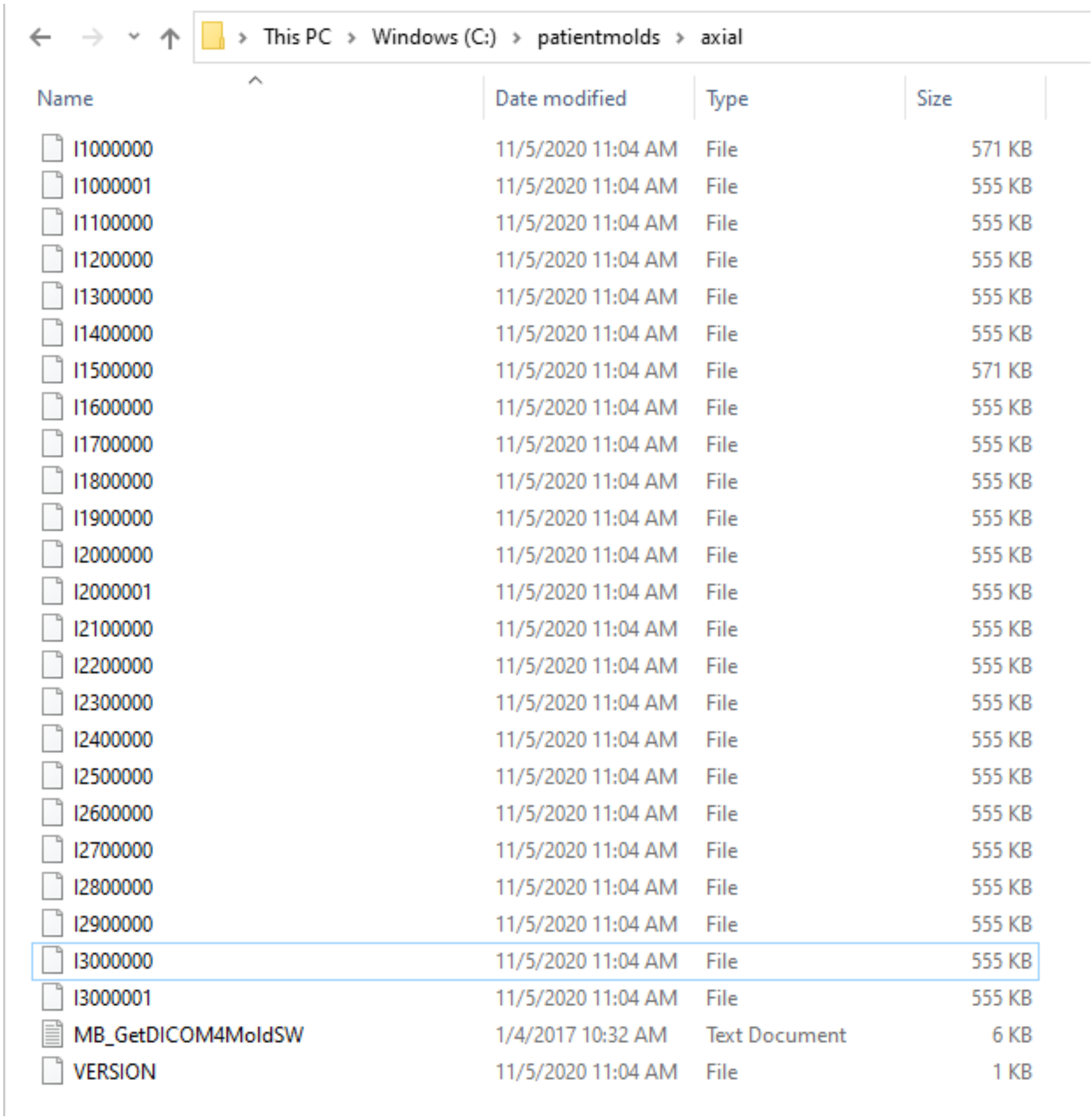

> This PC > Windows (C:) > patientmolds > axial

| Name | Date modified | Type | Size |
|---|---|---|---|
| I1000000 | 11/5/2020 11:04 AM | File | 571 KB |
| I1000001 | 11/5/2020 11:04 AM | File | 555 KB |
| I1100000 | 11/5/2020 11:04 AM | File | 555 KB |
| I1200000 | 11/5/2020 11:04 AM | File | 555 KB |
| I1300000 | 11/5/2020 11:04 AM | File | 555 KB |
| I1400000 | 11/5/2020 11:04 AM | File | 555 KB |
| I1500000 | 11/5/2020 11:04 AM | File | 571 KB |
| I1600000 | 11/5/2020 11:04 AM | File | 555 KB |
| I1700000 | 11/5/2020 11:04 AM | File | 555 KB |
| I1800000 | 11/5/2020 11:04 AM | File | 555 KB |
| I1900000 | 11/5/2020 11:04 AM | File | 555 KB |
| I2000000 | 11/5/2020 11:04 AM | File | 555 KB |
| I2000001 | 11/5/2020 11:04 AM | File | 555 KB |
| I2100000 | 11/5/2020 11:04 AM | File | 555 KB |
| I2200000 | 11/5/2020 11:04 AM | File | 555 KB |
| I2300000 | 11/5/2020 11:04 AM | File | 555 KB |
| I2400000 | 11/5/2020 11:04 AM | File | 555 KB |
| I2500000 | 11/5/2020 11:04 AM | File | 555 KB |
| I2600000 | 11/5/2020 11:04 AM | File | 555 KB |
| I2700000 | 11/5/2020 11:04 AM | File | 555 KB |
| I2800000 | 11/5/2020 11:04 AM | File | 555 KB |
| I2900000 | 11/5/2020 11:04 AM | File | 555 KB |
| I3000000 | 11/5/2020 11:04 AM | File | 555 KB |
| I3000001 | 11/5/2020 11:04 AM | File | 555 KB |
| MB_GetDICOM4MoldSW | 1/4/2017 10:32 AM | Text Document | 6 KB |
| VERSION | 11/5/2020 11:04 AM | File | 1 KB |

**Step 4: Open the first DICOM image in the stack (e.g.: I1000000) using ImageJ.**

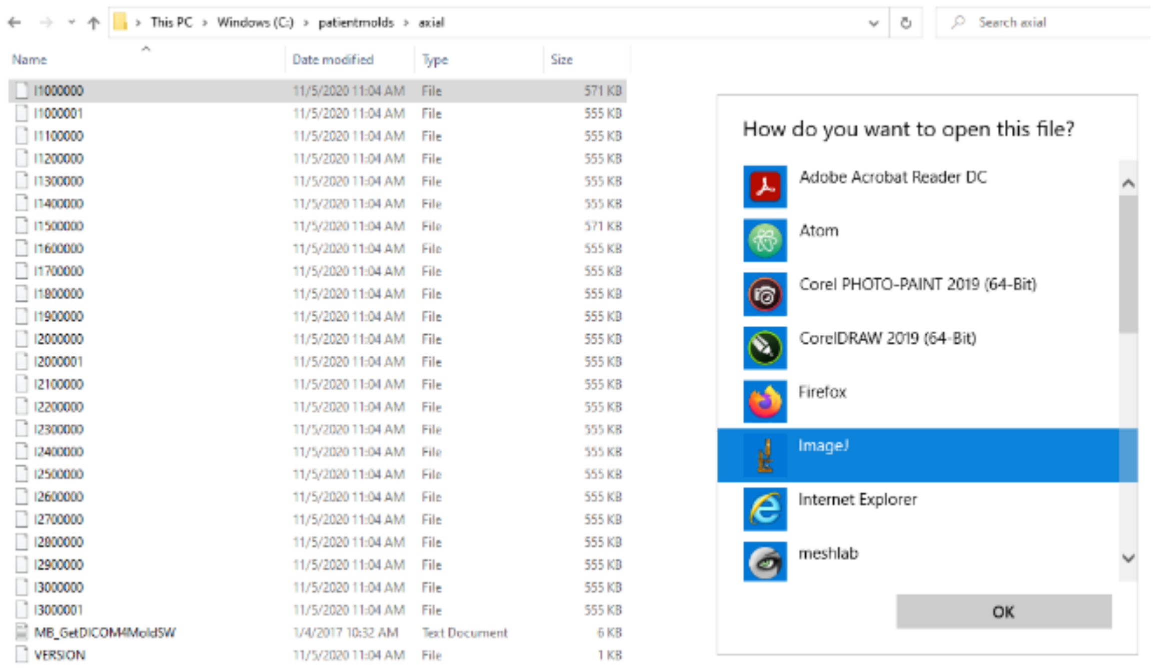


**Step 5: In the ImageJ menu, go to Plugins>Macros>Run, and select the "MB_GetDICOM4MoldSW" script.** A file named "dicom.txt" will be created in the "C:\patientmolds\mold" folder. An error message will occur if open DICOM file is not the first slice.

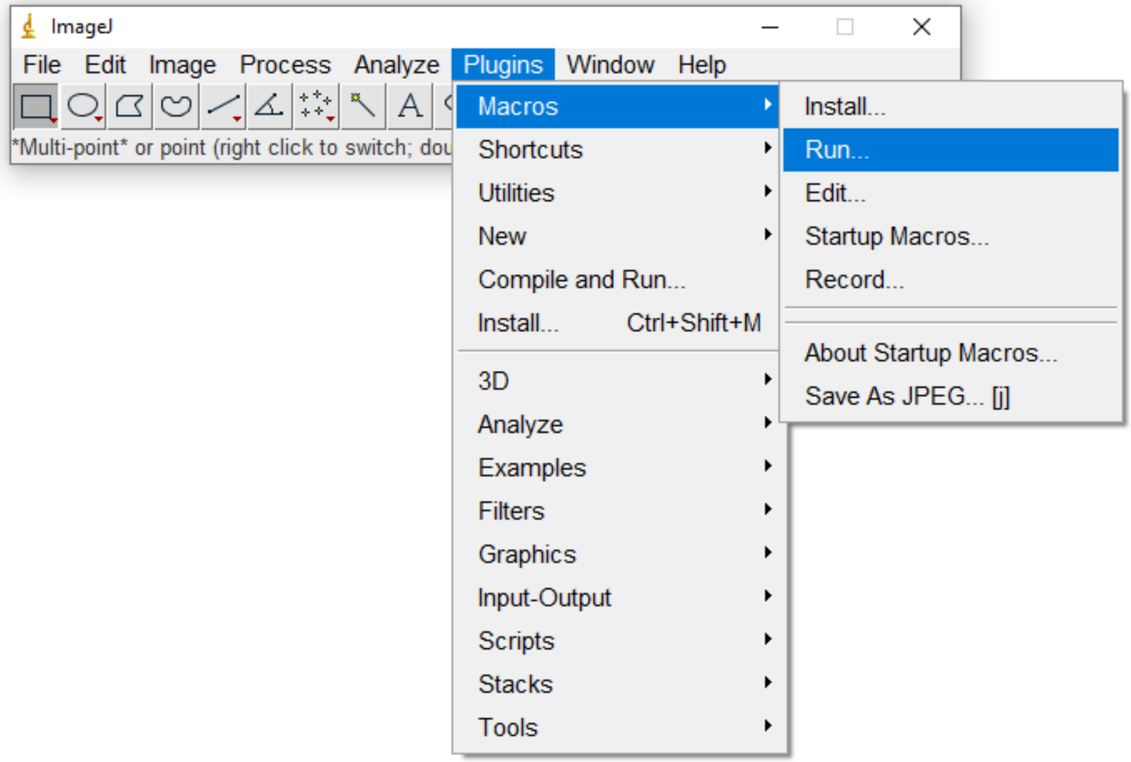


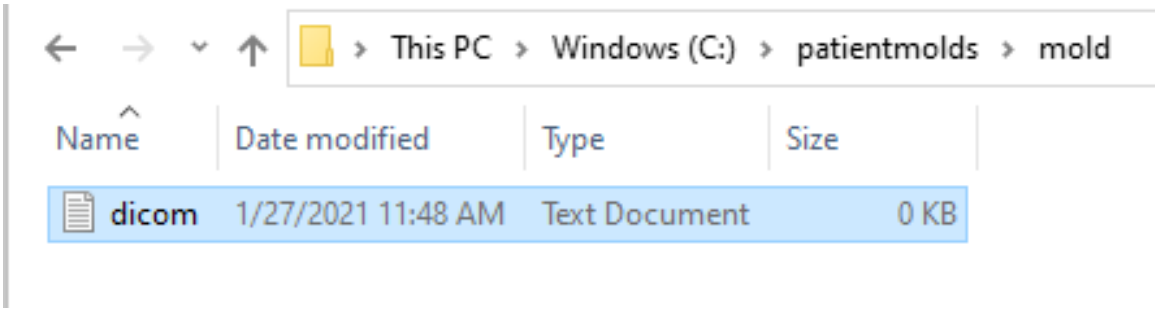


1.2 <u>VOI input</u>:

Required file: prostate axial VOI ("axial_final.voi")
Optional files: lesion and urethra VOIs ("tumor.voi", "urethra.voi"). If not provided, a prompt window will occur to confirm it.

Note: The macro requires specified names for the VOI files. However, these names can be modified within the code.

**Step 1: Create the "voi" folder in the "C:\patientmolds\mold" folder.**

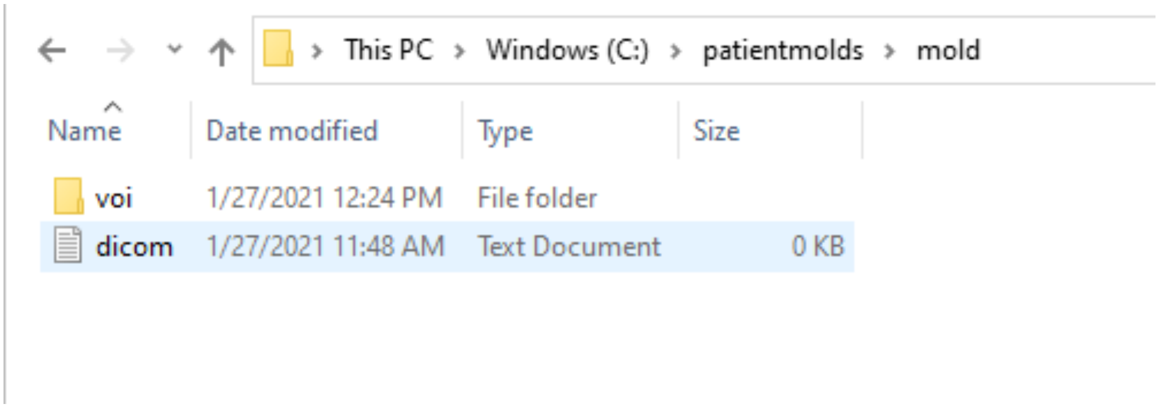


**Step 2: Place your .VOI files in the "C:\patientmolds\mold\voi" folder.**

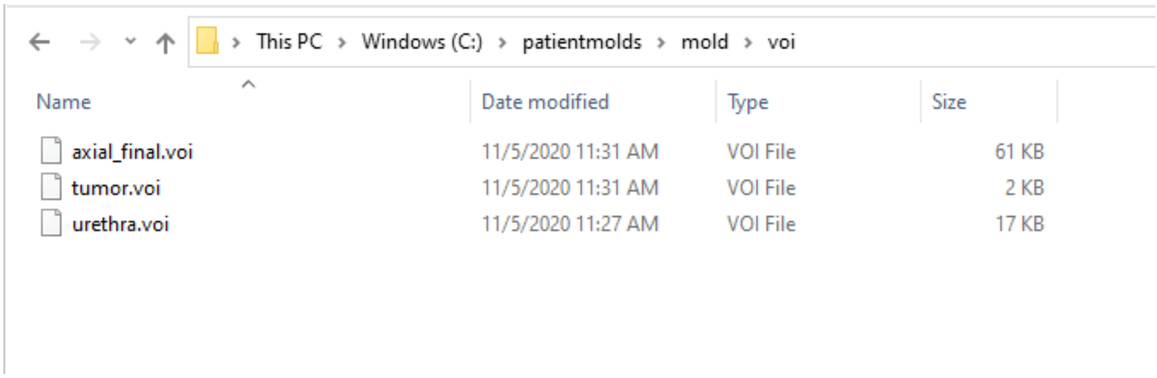


1.3 STL input:

Due to SolidWorks limitation, the STL files should not exceed 9000 faces. Note: There are a number of surface reconstruction software capable of "decimating" the 3D model to the allowed number of faces, e.g., Meshlab's "Simplification: Quadratic Edge Collapse Decimation" filter. Note: Lesion and urethra STL files are optional.

**Step 1: Create the "stl" folder in the "C:\patientmolds\mold" folder.**

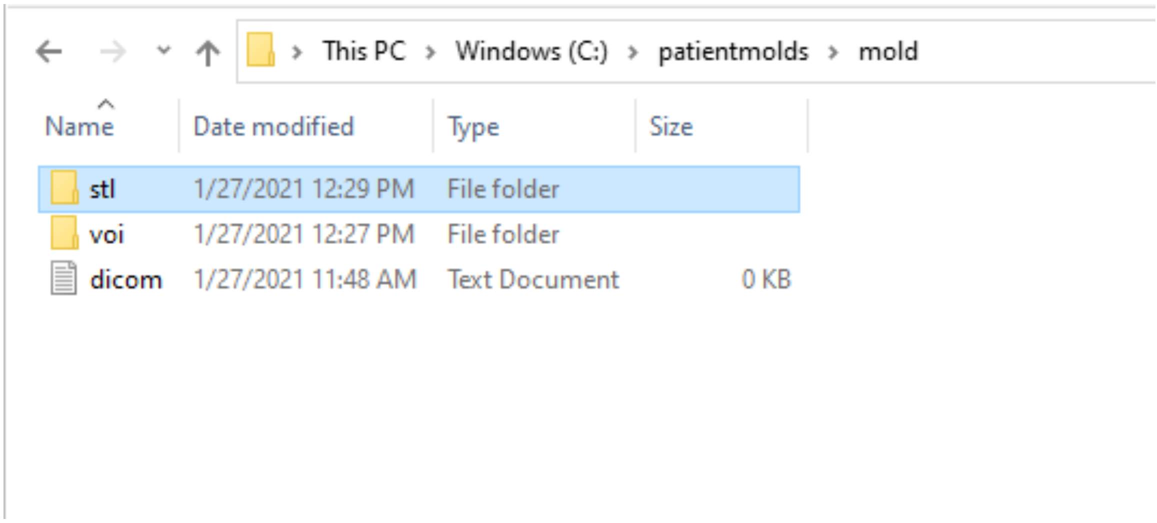


**Step 2: Place your .VOI files in the "C:\patientmolds\mold\stl" folder.**

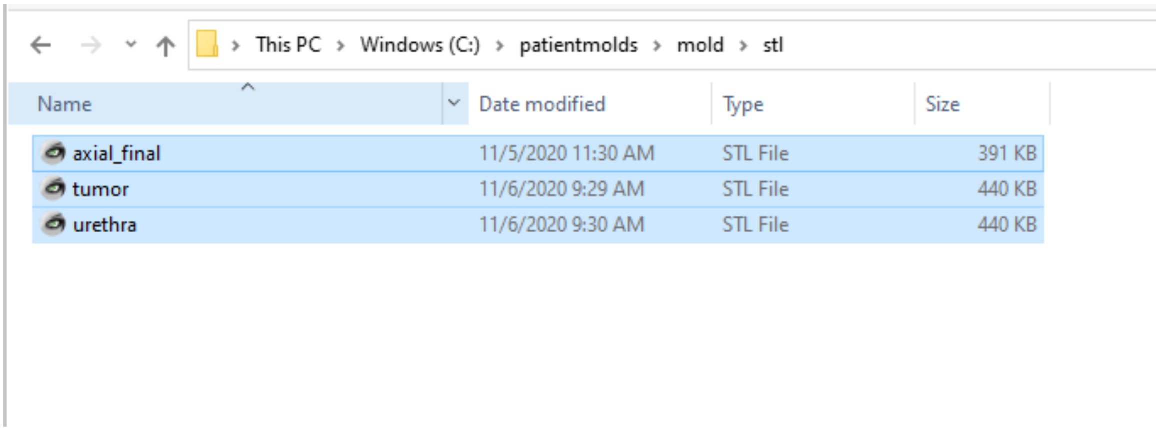


### 2. SolidWorks Macro:

Required software: SolidWorks 2020*
SolidWorks macro: “prostateMold_automationScript.swp”
SolidWorks settings: “prostateMold_swSettings_2020.sldreg”.

The file “prostateMold_swSettings_2020.sldreg” contains SolidWorks settings required to run the macro and is imported using the “Copy Settings Wizard”. Otherwise, SolidWorks should be configured with the following settings:

- Import STL as solid body
- Disable feature recognition on imports
- Units are set to millimeters

*Different versions of SolidWorks may not work properly. The SW macro should be edited to correctly reference the Part and Assembly template locations (e.g. for SW2018: C:\ProgramData\SOLIDWORKS\SOLIDWORKS 2018\templates).

**Step 1: Open SolidWorks 2020, and go to Tools>Macro>edit, and open “prostateMold_automationScript.swp”.**

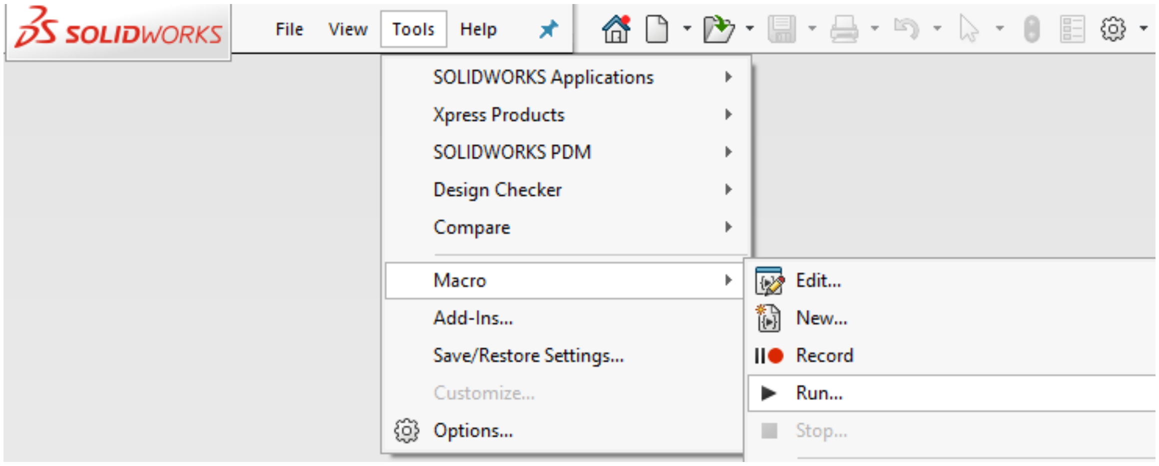

**Step 2: From the menu on the left, choose "verified_process1" inside the "Modules" folder by double clicking it.**

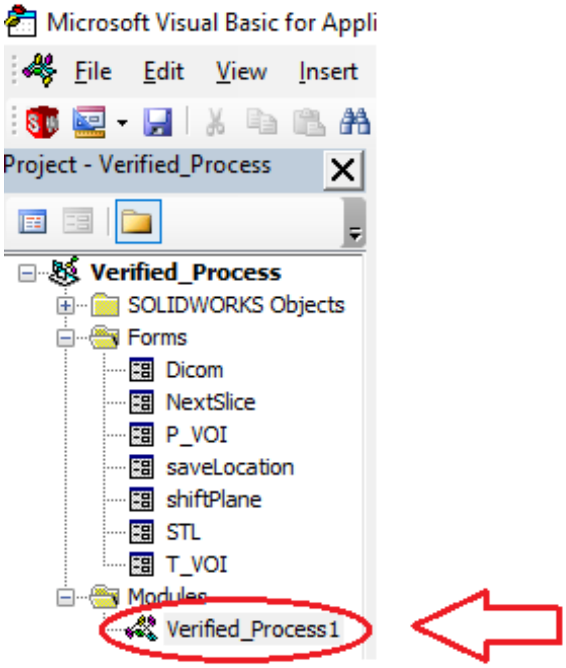


**Step 3: On the top menu, go to Run>Run sub/userform, or click on the green play button.**

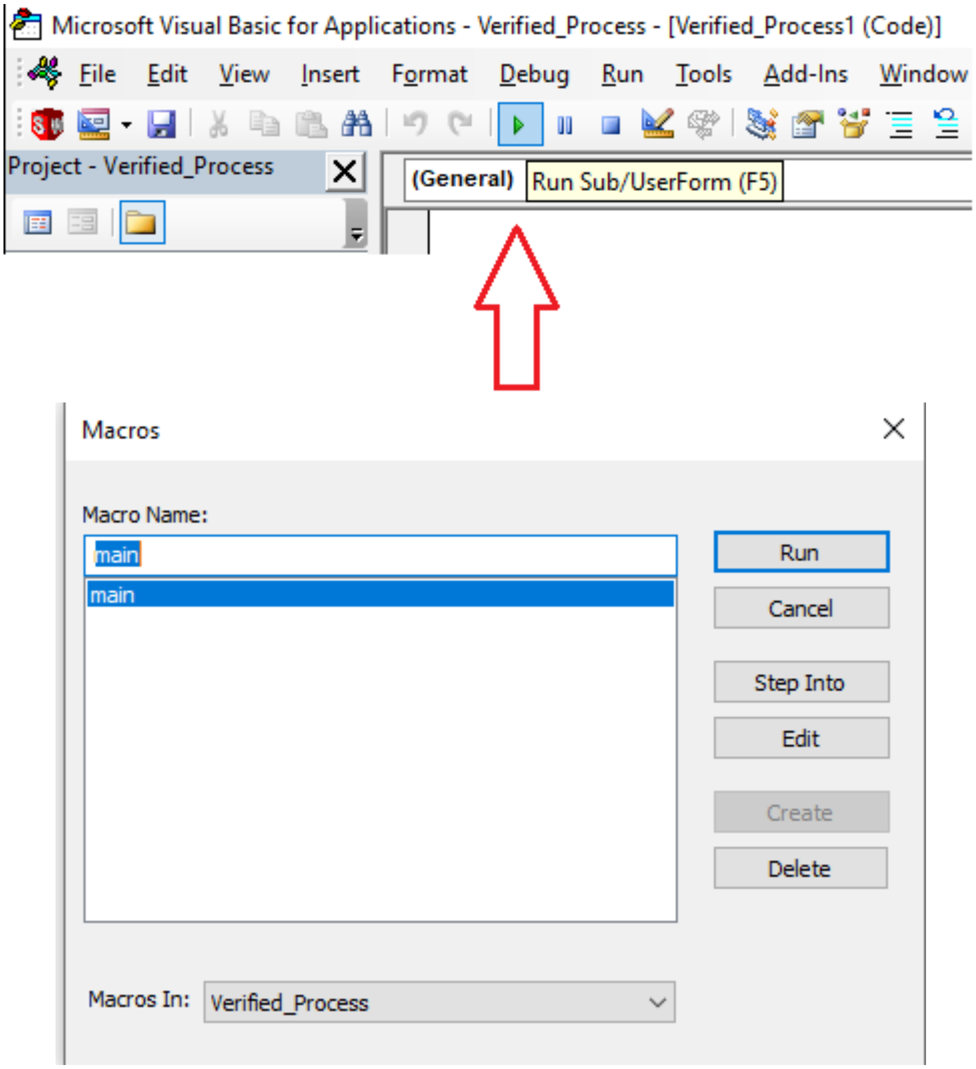


**Step 4: Click on Browse and select the prostate .STL file found in the "C:\patientmolds\mold\stl" folder.**

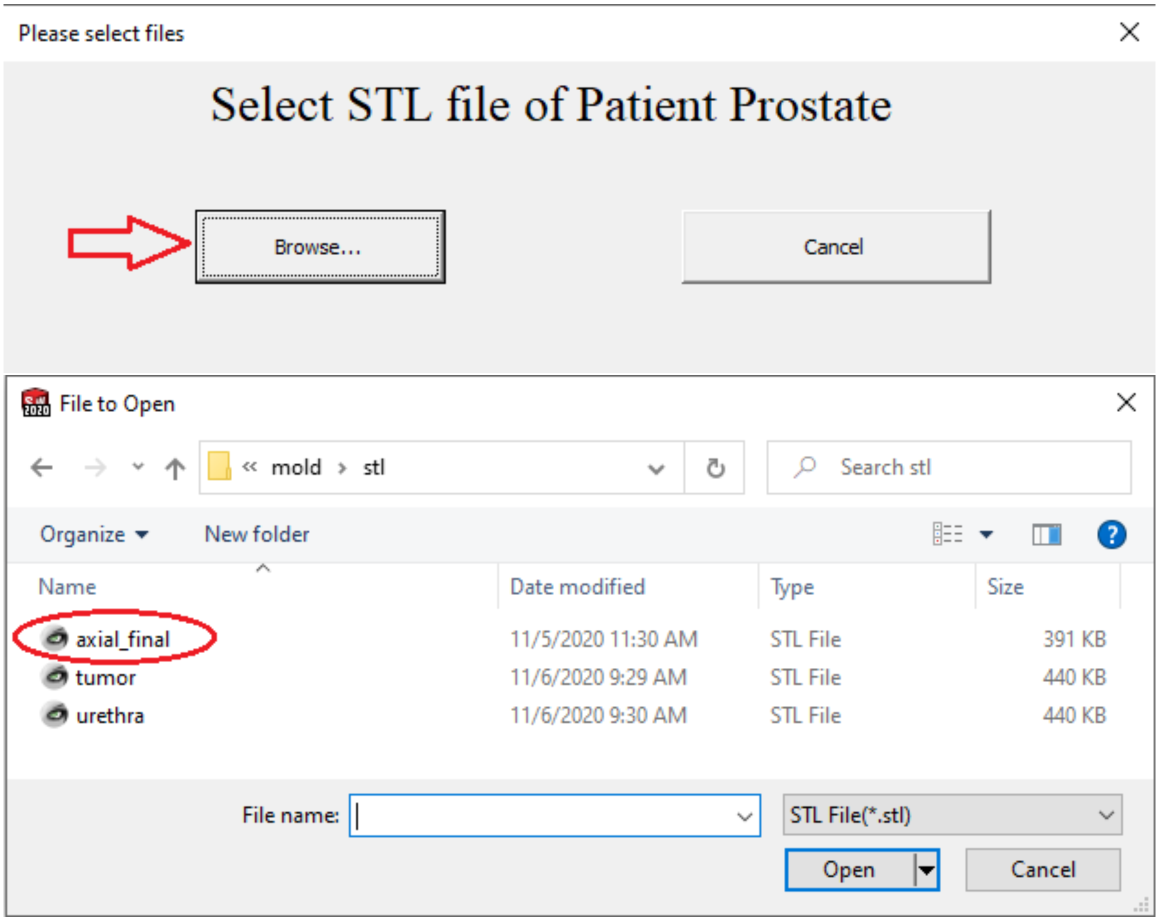


**Step 5: Run the macro. It may take a few minutes to complete.**

3. **Outputs:**

The macro-generated “C:\patientmolds\mold\output” folder contains the 2-piece mold, the locator card, and the 3D PDF assembly files. The current date is incorporated as a prefix in the file names.

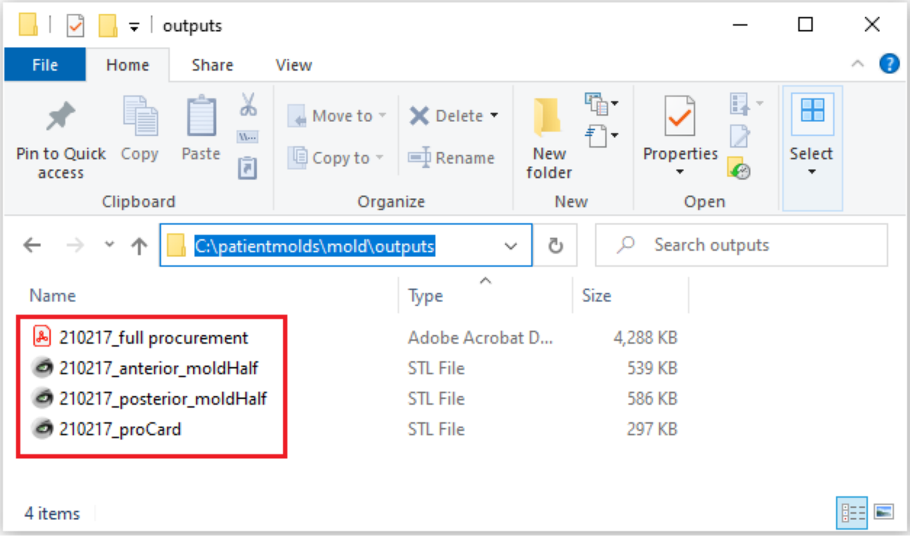
